# An unexpected donor in the adaptive introgression candidate *Helianthus annuus* subsp. *texanus*

**DOI:** 10.1101/2020.11.26.398933

**Authors:** Gregory. L Owens, Marco Todesco, Natalia Bercovich, Jean-Sébastien Légaré, Nora Mitchell, Kenneth D. Whitney, Loren H. Rieseberg

## Abstract

Hybridization is widely acknowledged as an important mechanism of acquiring adaptive variation. In Texas, the sunflower *Helianthus annuus* subsp. *texanus* is thought to have acquired herbivore resistance and morphological traits via introgression from a local congener, *H. debilis*. Here we test this hypothesis using whole genome sequencing data from across the entire range of *H. annuus* and possible donor species, as well as phenotypic data from a common garden study. We find that although it is morphologically convergent with *H. debilis, H. a. texanus* has conflicting signals of introgression. Genome wide tests (Patterson’s D and TreeMix) only find evidence of introgression from *H. argophyllus* (sister species to *H. annuus* and also sympatric), but not *H. debilis*, with the exception of one individual of 109 analysed. We further scanned the genome for localized signals of introgression using PCAdmix and found minimal but non-zero introgression from *H. debilis* and significant introgression from *H. argophyllus*. Putative introgressions mainly occur in high recombination regions as predicted by theory if introgressed ancestry contains maladaptive alleles. To reconcile the disparate findings of our analyses, we discuss potential test-specific confounding features, including introgression from other taxa. Given the paucity of introgression from *H. debilis*, we argue that the morphological convergence observed in Texas is likely independent of introgression.

## Introduction

Introgression is now widely appreciated to be an important force in evolution, as it is associated with many fundamental processes including speciation, adaptive radiation and invasion (Seehausen 2004; Abbott et al., 2013; Hovick & Whitney 2014; Arnold & Kunte 2017). Speciation and adaptation can occur very rapidly and in some cases the alleles responsible for these processes are derived from other species (e.g. Jay et al., 2018; Giska et al., 2019). For example, brown winter coats in snowshoe hares likely originated through introgression with black-tailed jackrabbits (Jones et al., 2018). In this case, the introgression is adaptive, since evidence suggests selection favored the introgressed allele. It is likely, however, that most introgression is not adaptive, but rather neutral or even maladaptive. Hybridization also combines speciation genes or incompatibilities, and empirical evidence supports that introgressed ancestry is often selected against (Harrison & Larson 2014). To better understand the overall role of introgression we need to better understand the rarer cases of adaptive introgression.

Sunflowers are a particularly well suited system for studying hybridization and introgression. Hybrids can be formed between most species in the annual clade, although strong sterility barriers exist in the F1 (Owens & Rieseberg 2015). The common sunflower, *Helianthus annuus*, and its sympatric congener, the prairie sunflower *H. petiolaris*, are known to hybridize and exchange alleles. This is seen both in the presence of hybrid swarms, and by model-based methods to estimate gene flow (Heiser 1947; Yatabe et al., 2007). It is thought that ancient hybridization between these species produced three different hybrid species, *H. anomalus*, *H. deserticola* and *H. paradoxus* (Rieseberg et al., 1990b, Rieseberg 1991). The common sunflower has a wide range across the United States, as well as southern Canada and northern Mexico, and it has more recently spread into California, where it absorbed alleles from the native *H. bolanderi* (Heiser 1949; Owens et al., 2016). It is possible that the wide range of *H. annuus* is due to its ability to incorporate locally adaptive alleles from congeners.

This hypothesis is best highlighted by studies of Texas populations. Populations of *H. annuus* in Texas have been found to be morphologically convergent with a local congener, *H. debilis* var. *cucumerifolius*, at several traits including purple mottled stems, smaller flower heads and earlier flowering (Heiser 1951a). This, combined with observations that *H. annuus* was exclusive to human disturbed locations, and documented hybridization between the two species, led Heiser to argue that *H. annuus* recently expanded into eastern central Texas and that introgression from *H. debilis* facilitated it (Heiser 1951a; Heiser 1951b). Since populations of *H. annuus* are interfertile across their range, these putatively introgressed populations in Texas were considered ecotypes and later given subspecific status as *H. annuus* subsp. *texanus* (Heiser 1954), while populations in the rest of the range are referred to as *H. annuus* subsp. *annuus*.

Another sunflower species, the silverleaf sunflower *H. argophylllus*, is also found in Texas, but has generally not been thought to have contributed to the formation of *H. a. texanus* (Heiser 1954). Although Heiser recognized that *H. argophyllus* was a possible donor due to its location and the presence of hybrids with *H. annuus*, *H. debilis* seemed a more likely choice, since it shares several distinctive traits with *H. a. texanus* (Heiser 1951b). In contrast, while *H. argophyllus* has relatively small flower heads, similar to *H. a. texanus*, it does not have other morphological characteristics that are typical of this subspecies, like visible purple mottled stems, and generally flowers much later than *H. annuus*, rather than earlier. The silverleaf sunflower is also characterized by, and named for, the dense white pubsence on its stem and leaves, a trait not found in *H. a. texanus*. Additionally, hybrids between the species had strong F1 sterility, limiting the likelihood of introgression (Chandler et al., 1986).

More recently, molecular analyses have tested this hypothesis using rDNA, cpDNA, AFLPs and microsatellites, each finding evidence that alleles diagnostic for *H. debilis* can be found in *H. a. texanus* at low frequency (Rieseberg et al., 1990a, 2007; Scascitelli et al., 2010). Furthermore, field experiments found that local subsp. *texanus* had higher fitness than subsp. *annuus* when grown in Texas, and that fitness-related trait values were shifted towards *H. debilis* (Whitney et al., 2006, 2010). In those experiments, the pattern of selection favoring local trait values was also seen in BC1 hybrids between subsp. *annuus* and *H. debilis*, supporting the adaptive potential of introgression. Finally, an eight-year field experimental evolution study found that *H. a. annuus* x *H. debilis* hybrids (synthesized to mimic the putative early ancestors of *H. a. texanus*) rapidly evolved higher fitness than non-hybrid *H. a. annuus* controls when exposed to the central Texas environment (Mitchell et al., 2019). Together these results suggest that *H. a. texanus* has introgressed ancestry from *H. debilis* and that these alleles should control adaptive and convergent morphological traits.

In this study, we compare common garden morphology measurements between *H. annuus* subspecies and use whole genome sequence data to examine the genome wide ancestry of *H. a. texanus*. Our main goal is to identify how much of the genome is derived from *H. debilis*, or from another congener, *H. argophyllus*, not previously suspected as being involved. Although *H. a. texanus* has been examined genetically before, this is the first analysis using modern genomics and methods that distinguish incomplete lineage sorting and introgression. We use these methods to quantify introgression at a full genome and genomic window level and we discuss how our results change our understanding of this story.

## Methods

### Morphological analysis

Before evaluating the genetics of *H. a. texanus*, we first re-assessed whether *H. annuus* samples from Texas were morphologically distinct from samples from the rest of the species range. The original subspecies designation by Heiser (1954) suggested that *H. a. texanus* was found in eastern Texas, and graded into other subspecies in northern and western Texas. Taking this observation into account, we divided our samples into “southern Texas” samples below 30° latitude, “northern Texas” samples above 30° latitude but within Texas, and “non-Texas”, including all others. We assessed the morphology using data collected from a previously reported common garden of *H. annuus* (Todesco et al., 2020). Briefly, 614 *H. annuus* from 63 populations across the native range were grown in Vancouver, BC, Canada and measured for a range of morphological and phenological traits.

Based on the original description of *H. a. texanus*, we examined traits that were expected to differ between subsp. *annuus* and subsp. *texanus*, and their expected direction. We then used R to conduct a Mann-Whitney test for each variable asking if the southern and/or northern Texas populations differed from the rest of the species (R core team, 2013). To see if Texas samples are exceptional in general, we ran a principal component analysis (PCA) of all traits using FactoMineR, and missing data imputed with MissMDA (Lê et al., 2008; Josse & Husson, 2016). Based on this analysis, when directly comparing *H. a. annuus* and *H. a. texanus* as a whole, we included southern Texas samples (identifying them as subsp. *texanus*) and the non-Texas samples (identifying them as subsp. *annuus*) and excluded intermediate north Texas samples. When analysing individual samples, we included all Texas *H. annuus* to probe possible geographic patterns. Figure colors were chosen from the PNWColors palettes (Lawlor, 2020)

### Population genetics

From a previously published whole genome resequencing dataset, we selected all *H. annuus* samples from Texas, and one random sample per population for the remaining wild *H. annuus*, as well as from the annual species *H. argophyllus*, *H. debilis*, *H. petiolaris* subsp. *fallax*, *H. petiolaris* subsp. *petiolaris* and *H. niveus* subsp. *canescens* (Todesco et al., 2020). We also selected four outgroup perennial samples, one from each of *H. divaricatus*, *H. giganteus*, *H. decapetalus* and *H. grosseserratus*. All samples were variant-called together using the same pipeline as Todesco et al. (2020). All sample information, including SRA accession numbers for raw sequence data, are collated in File S1. Briefly, samples were trimmed using Trimmomatic, aligned to the *H. annuus* XRQv1 reference genome using NextGenMap, and variant-called using GATK. Variants were filtered using the GATK VQSR using a set of 67 cultivated *H. annuus* as the truth set, and the 90% tranche was retained (Bolger et al., 2014; Badouin et al., 2017; Sedlazeck et al., 2013; Van der Auwera et al., 2013). The dataset was further filtered to retain only bi-allelic SNPs with minor allele frequency ≥ 1% and a genotyping rate ≥ 90%.

Variants were then remapped to the *H. annuus* Ha412HOv2 using BWA, as this has been shown to dramatically improve SNP ordering (Todesco et al., 2020). Finally, the dataset was phased and imputed using Beagle (Browning & Browning, 2007). The dataset was subset for further analyses using bcftools and converted to specific program input formats using plink and custom perl scripts (Li et al., 2011; Purcell et al., 2007). All scripts are available at https://github.com/owensgl/texanus_ancestry.

We calculated genome-wide Weir and Cockerham F_ST_ between *H. a. texanus*, *H. a. annuus*, *H. argophyllus* and *H. debilis* using a custom perl script (Weir & Cockerham, 1984), requiring a minor allele frequency ≥ 1% for each locus in tested samples. Our sampling strategy was unbalanced and included fewer samples but more populations for subsp. *annuus* compared to subsp. *texanus* (59 samples in 59 populations vs 109 samples in 11 populations), and this combined with minor allele frequency filters may bias results (i.e. if rare population-specific alleles are being preferentially retained in one set). To combat this, when comparing F_ST_ scores between subsp. *annuus* and subsp. *texanus*, we filtered loci so that both comparisons had an F_ST_ value (that is, the site is variable and above MAF cut offs in both *H. annuus* groups).

Additionally, we subsampled down to 11 samples from different populations for each *H. annuus* group, required a minor allele frequency ≥ 5% and repeated F_ST_ calculations. We then visualized these results in 1 Mbp non-overlapping sliding windows, with at least 10 loci, by summing the numerator and denominator of F_ST_ within the window, using the tidyverse library in R (Wickham et al., 2019). We tested if there was a significant difference in window F_ST_ between the *H. annuus* groups and *H. argophyllus* or *H. debilis* using a paired T-test. Additionally, we imputed the recombination rate for each F_ST_ window using a consensus genetic map built from five domestic *H. annuus* genetic maps (Todesco et al., 2020). We visualized F_ST_ in five quantiles divided by ranked recombination rate.

### Genome wide introgression analysis

To detect admixture, we calculated Patterson’s D using Admixr, an R wrapper for Admixtools (Patterson et al., 2012; Petr et al., 2019). D, also known as the ABBA-BABA test, asks if there is a greater number of shared derived alleles between taxa than expected under incomplete lineage sorting. In this case, we ask if *H. a. texanus* shares more derived alleles with a donor species (e.g. *H. debilis*) than *H. a. annuus*, which is allopatric and should not be affected by gene flow (although see below). We used two possible donor species: *H. debilis*, the hypothesized donor, and *H. argophyllus*.

One possible confounding feature of D is when there are multiple hybridization events in different parts of the tree, the signal of specific introgression events can be masked. Previous work has suggested ongoing gene flow between *H. annuus* and *H. petiolaris*, and we expect that this most recently occurs in the non-Texas populations of *H. a. annuus* since *H. petiolaris* has a limited range in Texas. Since *H. debilis* is closely related to *H. petiolaris*, introgression between *H. a. annuus* and *H. petiolaris* may mask introgression from *H. a. texanus* and *H. debilis* (Fig. 3B). To explore this possible issue, we also used *H. petiolaris fallax* as a possible donor species.

For each *H. a. texanus* sample, we calculated D using the arrangement [a single *H. a. texanus* sample, all *H. a. annuus*, potential donor, all perennials]. We ran this independently using *H. argophyllus*, *H. debilis or H. petiolaris fallax* as the potential donor in position 3. We used the default (1 Mbp) window size for block bootstrapping. This allowed us to look for variation in introgression between individuals to determine if introgression is present in all *H. a. texanus* samples. We also ran the following arrangements to test for *H. argophyllus* - *H. debilis* introgression using the following sets: [*H. debilis*, *H. petiolaris fallax*, *H. argophyllus,* perennials], [*H. a. texanus*, *H. argophyllus*, *H. debilis*, perennials], [*H. debilis*, *H. petiolaris fallax*, *H. a. texanus*, perennials] and [*H. debilis*, *H. petiolaris fallax*, *H. a. annuus*, perennials]. For tests comparing whole species, rather than populations, we repeated these tests using *H. petiolaris petiolaris* instead of *H. petiolaris fallax* to check for subspecies effects.

Genomic admixture can also be detected using programs for identifying population structure, like ADMIXTURE (Alexander & Lange, 2011). To remove linkage, we used SNPrelate to prune for markers with high linkage (r^2^ ≥ 0.2 within 500 Kbp) and selected all *H. annuus*, *H. argophyllus* and *H. debilis* samples from our larger dataset (Zheng et al., 2012). We ran ADMIXTURE with K from 1 to 10 with 200 bootstrap replicates, and cross validation to determine the best fit K. We then used CLUMPAK to synchronize groupings between K values (Kopelman et al., 2012). We presented results at K = 5, which was the minimum K value that best separated known species. To confirm ADMIXTURE results, we ran STRUCTURE with K = 5 for 20 replicates each randomly subset to 5% of the total sites due to computational limitations (Pritchard et al., 2000). These results were also synchronized with CLUMPAK and we report the two major output patterns.

Another way of looking at gene flow between species is through the covariance of allele frequencies, which can be produced through shared drift or gene flow. TreeMix creates a tree of populations based on shared genetic drift and adds migration events to explain exceptional shared drift (Pickrell & Pritchard, 2012). This method requires unlinked markers, so we used SNPrelate to prune out markers with high linkage as above. For this analysis, we included all species in our dataset and divided *H. annuus* into three groups: non-Texas (*H. a. annuus*), south Texas (*H. a. texanus*), and north Texas (intermediate). We also included *H. niveus canescen*s to better represent the full annual clade. Each tree was rooted with the four perennial species which were grouped into a single population for simplicity. We plotted the residual fit from the maximum likelihood tree by dividing the residual covariance between pairs of populations by the average standard error across all pairs. We selected an appropriate number of migration edges based on the decay in likelihood improvements with successive numbers.

Since the initial TreeMix analyses did not include a migration branch from *H. debilis* into south Texas *H. annuus*, we specifically tested the likelihood of migration from *H. debilis* or from *H. argophyllus*. For each potential donor, we manually added a single migration event, with to 0.4 admixture proportion, and recorded the likelihood. We plot the likelihood relative to the highest likelihood proportion.

### Introgression candidates

To identify potentially introgressed regions at each site in the genome we measured D and *f_d_* using a custom perl script (Martin et al., 2014). This approach uses allele frequencies instead of genotypes from individual samples, and allows us to extract window values. We visualized this in 100 SNP non-overlapping windows and selected the top 1% of windows based on *f_d_* as outliers and potential introgressed regions. We used *f_d_* instead of D because it is more robust to low diversity regions (Martin et al., 2014). We also ran this analysis for a single *H. a. texanus* sample (ANN1363) that was identified as having extra *H. debilis* ancestry in ADMIXTURE and genome wide D.

Another way of detecting introgression is using genomic regions where samples do not genetically cluster with others of their species. We applied this approach using PCAdmix (Brisbin et al., 2012). In this analysis we included all samples from *H. a. annuus*, *H. argophyllus* and *H. debilis* as parental populations and all *H. a. texanus* samples as potentially admixed. Genetic map positions were imputed from a 1 Mbp resolution genetic map for Ha412HOv2 (Todesco et al., 2020). In regions with zero recombination across 1 Mbp, we smoothed the cM position across the nearest neighbouring positions with a non-zero cM/Mbp rate. We filtered for linkage and used 100 SNP windows (−r^2^ 0.8 -w 100). We selected windows with *H. debilis* ancestry in > 40% of chromosomes as being candidate introgression regions. As a control, we reran the analysis pulling out a single allopatric *H. a. annuus* sample and testing it as an admixed sample, repeating for each *H. a. annuus* sample.

For both D and PCAdmix, we explored the role of recombination by plotting the D scores or introgression frequency in windows grouped by recombination rate quantile.

## Results

### Morphology

Consistent with the existing subspecies description (Heiser, 1954), *H. annuus* samples from south Texas tend to begin branching closer to the ground, have smaller inflorescences (measured as the diameter of the central disk), more purplish stem coloration, and smaller seeds (Fig. 1C-F), although their values are within the range found across the range of non-Texas *H. annuus*. Other trait values are also consistent with the subspecies description, including phyllary length and ligule number (Fig. S1). This morphological distinctiveness is seen in the trait PCA, which locates Texas samples along a restricted portion of the first PC axis, while non-Texas samples span the entire axis (Fig. 1A-B). North Texas samples are often morphologically intermediate (Fig. 1B), so moving forward we excluded these samples when testing *H. a. Texanus* as a whole.

**Figure 1:**
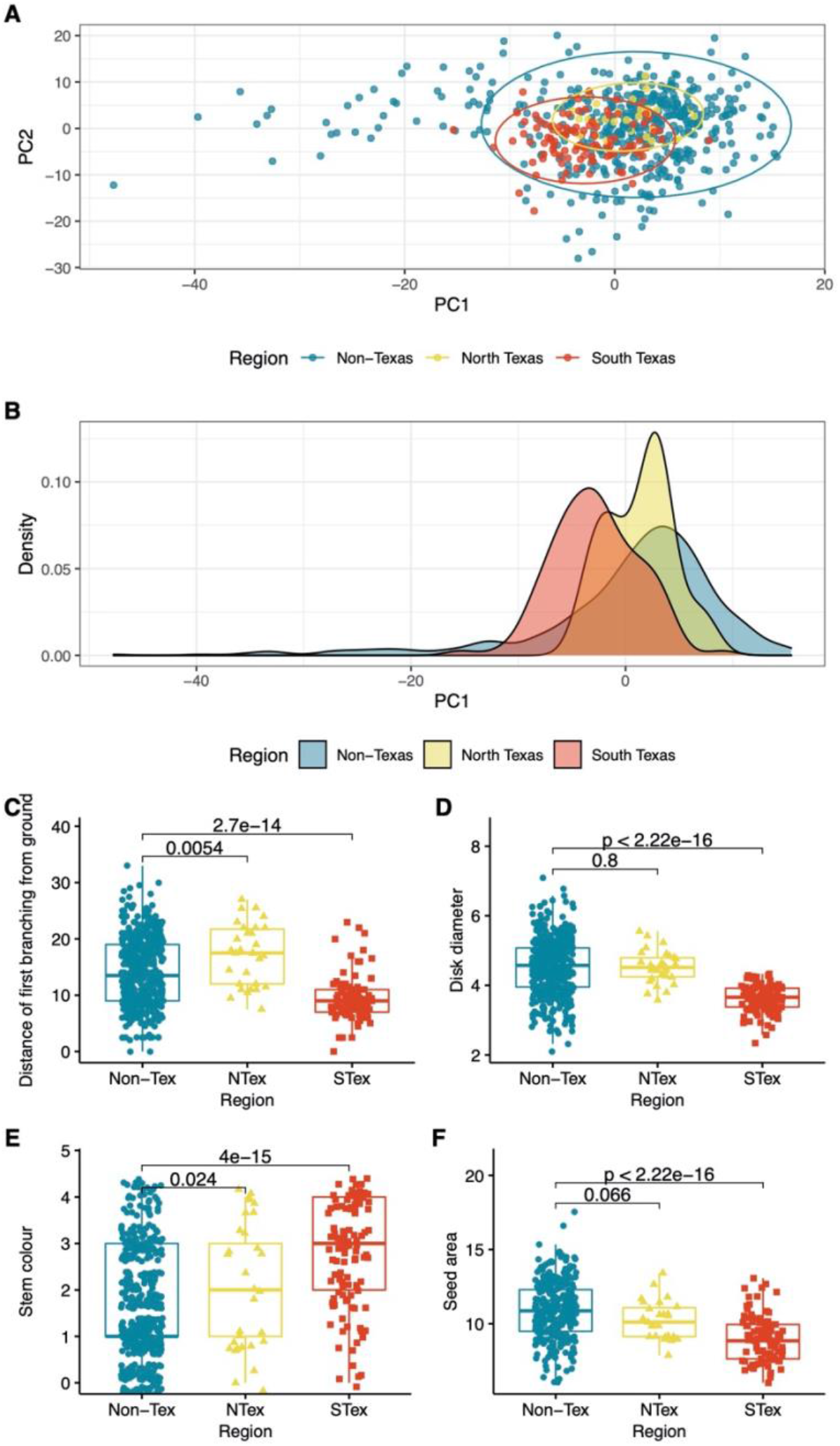
Phenotype PCA of H. annuus samples. A. The first two principal components for a PCA of H. annuus traits measured in a common garden. The ellipses cover 95% of the points in each group. B. A density plot of the first principal component divided by geographic region. C-F. Selected morphological traits known to differentiate H. a. annuus from H. a. texanus. P-values represent Wilcox rank sum tests against non-Texas H. annuus. For E, darker, more purple stems were represented by higher values.

### Population genetics

We found that *H. a. texanus* populations were moderately differentiated from *H. a. annuus* populations (F_ST_ = 0.0635; 0.0032-0.712 in 1 Mbp windows). Regions of high differentiation often occurred in previously identified putative segregating structural variants (“haploblocks”, Fig. 2A; Todesco et al., 2020). For both subspecies of *H. annuus*, there was high differentiation from *H. argophyllus* and *H. debilis*. F_ST_ was significantly lower for *H. a. texanus* samples than for *H. a. annuus* samples when both were compared to *H. argophyllus* 1 Mbp windows; *H. argophyllus* [paired t(3093) =30.923, p ≤ 2.2e-16], but only marginally lower when compared to *H. debilis* [paired t(3082) =1.7514, p = 0.08)] (Fig. 2B-D). When tested using a subset of samples, so both *H. annuus* subspecies had equal numbers of populations and samples, we obtained qualitatively similar results, although less differentiation in the *H. debilis* comparison; *H. argophyllus* [paired t(3075) =18.871, p ≤ 2.2e-16] and *H. debilis* [paired t(3074) =-0.5312, p = 0.5)]. We further found that the lowest F_ST_ is found in regions of high recombination in both comparisons (Fig. 2E,F).

**Figure 2:**
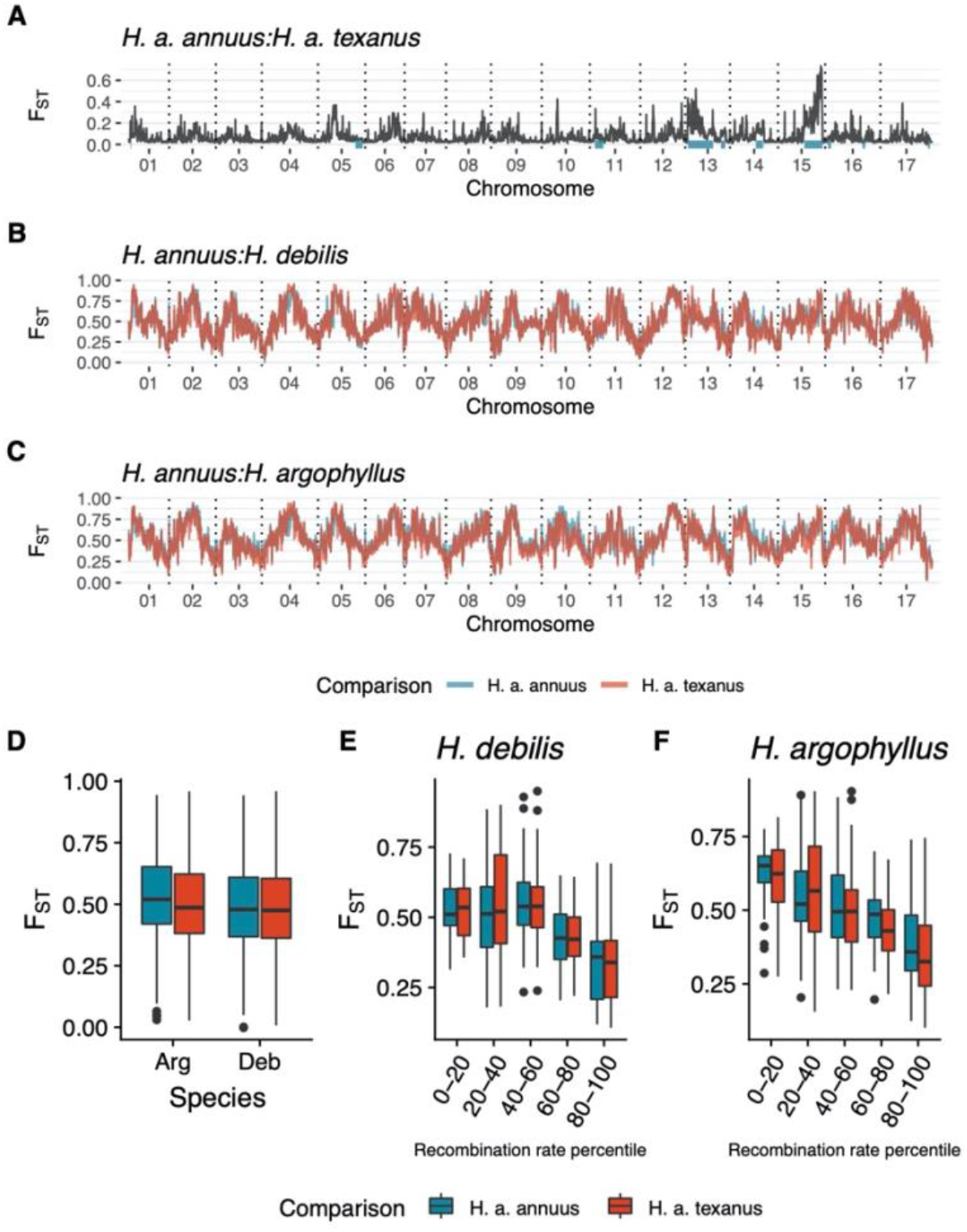
F_ST_ between H. a. annuus, H. a. texanus and congeners. A. F_ST_ between H. a. annuus and H. a. texanus in 1 Mbp windows. Genomic locations for previously identified haploblocks are highlighted in blue. B. F_ST_ between H. annuus and H. argophyllus in 1 Mbp windows. C. F_ST_ between H. annuus and H. debilis in 1 Mbp windows. D. Boxplot of F_ST_ in 1 Mbp windows. E-F. Comparison of F_ST_ with recombination rate based on a H. annuus genetic map. F_ST_ windows are binned into 20% percentile groups based on recombination rate.

### Introgression detection via ABBA-BABA tests

We calculated Patterson’s D for each Texas *H. annuus* sample, looking for introgression from *H. argophyllus*, *H. debilis* and *H. petiolaris fallax*. A majority of Texas *H. annuus* samples had a significant signal of introgression from *H. argophyllus*, and particularly high values from three populations, ANN_55, ANN_56 and ANN_57 (Fig. 3C). In contrast, a majority of samples had negative scores for *H. debilis* introgression (suggesting greater *H. debilis* allele sharing with allopatric *H. a. annuus* than with *H. a. texanus*). Higher *H. debilis* D values occurred in samples with high *H. argophyllus* D scores; *H. argophyllus* D significantly explained *H. debilis* D (R^2^ = 0.6585, p < e^−15^) (Fig. 3E).

**Figure 3:**
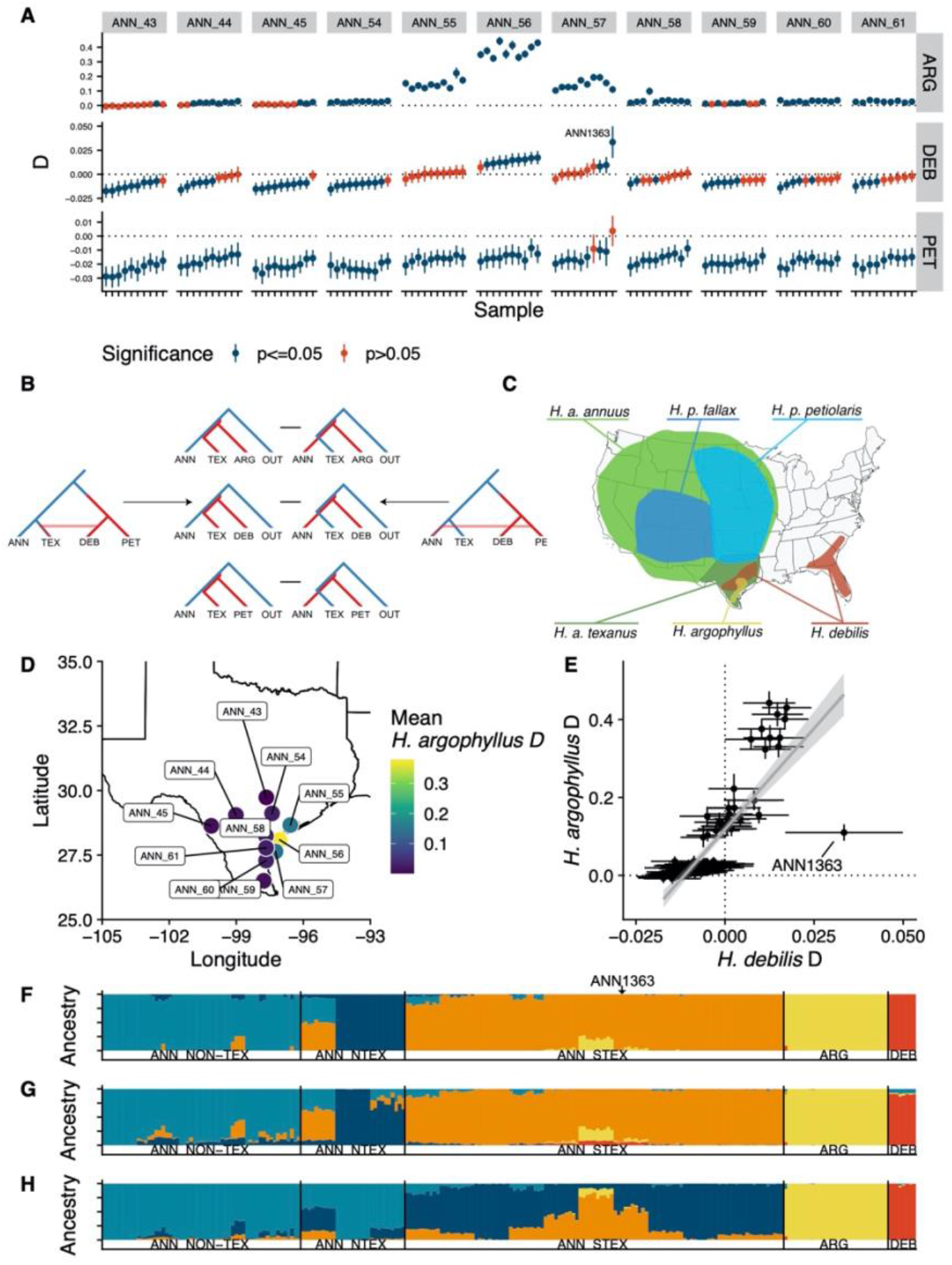
Measures of admixture in H. a. texanus. A. Patterson’s D with 95% bootstrap confidence intervals for Texas H. annuus samples with three possible donor species. Blue bars indicate haploblocks discovered in Todesco et al. ARG: H. argophyllus, DEB: H. debilis, PET: H. petiolaris fallax. B. Phylogenetic representation of D test for ARG, DEB and PET donors. Side trees represent how different introgression events could affect D tests involving DEB. C. The ranges of annual sunflowers used in this analysis. D. Mean Patterson’s D for Texas H. annuus populations with H. argophyllus as the possible donor. E. The relationship between Patterson’s D for H. a. texanus samples using H. argophyllus and H. debilis as donors. Error bars represent 95% bootstrap confidence interval. F. ADMIXTURE results for H. annuus, H. argophyllus and H. debilis samples with K=5. G. STRUCTURE results for 11/20 replicates. H. STRUCTURE results for 9/20 replicates.

Lastly, almost all *H. a. texanus* samples had a negative *H. petiolaris fallax* D, suggesting greater allele sharing between the parapatric *H. a. annuus* and *H. petiolaris fallax* than between the latter and *H. a. texanus*. We did find one *H. a. texanus* sample, ANN1363, with positive *H. debilis* D not accompanied by higher *H. argophyllus* D (Fig. 3E). We calculated genome wide D using just this sample and found localized evidence of introgression on chromosome 3, suggesting this is an advanced generation backcross individual (Fig. S2).

We also tested for introgression among Texas annual sunflower species outside of the focal *H. a. texanus* system. We found that there was ongoing gene flow between *H. argophyllus* and *H. debilis* (D = −0.049, p < e^−21^), consistent with the admixture signals seen in ADMIXTURE (see below). Despite this, there is a greater signal of gene flow between the allopatric *H. petiolaris* and *H. argophyllus* than between *H. debilis* and *H. argophyllus* (*H. petiolaris fallax*: D = −0.062, p < e^−21^; *H. petiolaris petiolaris*: D = −0.092, p < e^−39^). This is also true when using *H. a. annuus* or *H. a. texanus* instead of *H. argophyllus*, suggesting that this may reflect known introgression between *H. annuus* and *H. petiolaris* (*H. petiolaris fallax*: D = - 0.09, p < e^−50^; D = −0.081, p < e^−43^; *H. petiolaris petiolaris*: D = - 0.122, p < e^−72^; D = −0.112, p < e^−64^) (Yatabe et al., 2007).

### Introgression detection via ADMIXTURE and STRUCTURE

ADMIXTURE analysis best matched species designations with five groups, one primarily in each of non-Texas *H. annuus*, north Texas *H. annuus*, south Texas *H. annuus*, *H. argophyllus* and *H. debilis*. At this number of groups we saw *H. argophyllus* introgression into three populations of *H. a. texanus* previously identified using D, as well as *H. debilis* introgression (~2%) into ANN1363 (Fig. 3F, Fig. S3). We did not see any additional introgression from *H. debilis* into *H. a. texanus*. In contrast to ADMIXTURE, STRUCTURE produced two nearly equally supported results. In one result, *H. debilis* was largely its own cluster but contained introgression from non-Texas *H. annuus*. This result suggested *H. debilis* introgression into three *H. a. texanus* populations that also contain *H. argophyllus* introgression. In the other result, there is no *H. debilis* introgression into *H. a. texanus* except for the previously identified ANN1363 (Fig. 3F-G).

### Introgression detection via TreeMix

We explored gene flow between multiple species using TreeMix. The likelihood improvements declined after five migration edges so we present that value (Fig. 4A; Fig. S5L). At five migration edges, TreeMix found evidence for gene flow from *H. petiolaris* into *H. debilis*, *H. petiolaris fallax* into *H. niveus canescens*, *H. debilis* into *H. argophyllus*, *H. argophyllus* into *H. a. texanus* and an ancestral node into *H. petiolaris fallax* (Fig. 4A). The overall tree is largely consistent with previously described phylogeny, except *H. debilis* is thought to be sister to *H. petiolaris*, likely explaining the strongest migration edge bringing those two together (Stephens et al., 2015). Even at ten migration edges, TreeMix does not suggest migration from *H. debilis* into *H. a. texanus* (Fig. S4). Treemix selects migration edges based on additional positive residual covariance after the initial tree but we see that the residual covariance with *H. debilis* and *H. a. texanus* is actually negative and less in *H. a. texanus* than in other *H. annuus* subspecies (Fig. 4B). When specifically tested, the most likely migration edge between *H. debilis* and *H. a. texanus* is 0. In contrast, the migration edge between *H. argophyllus* and *H. a. texanus* has the highest support at 0.19 (Fig. 4C).

**Figure 4:**
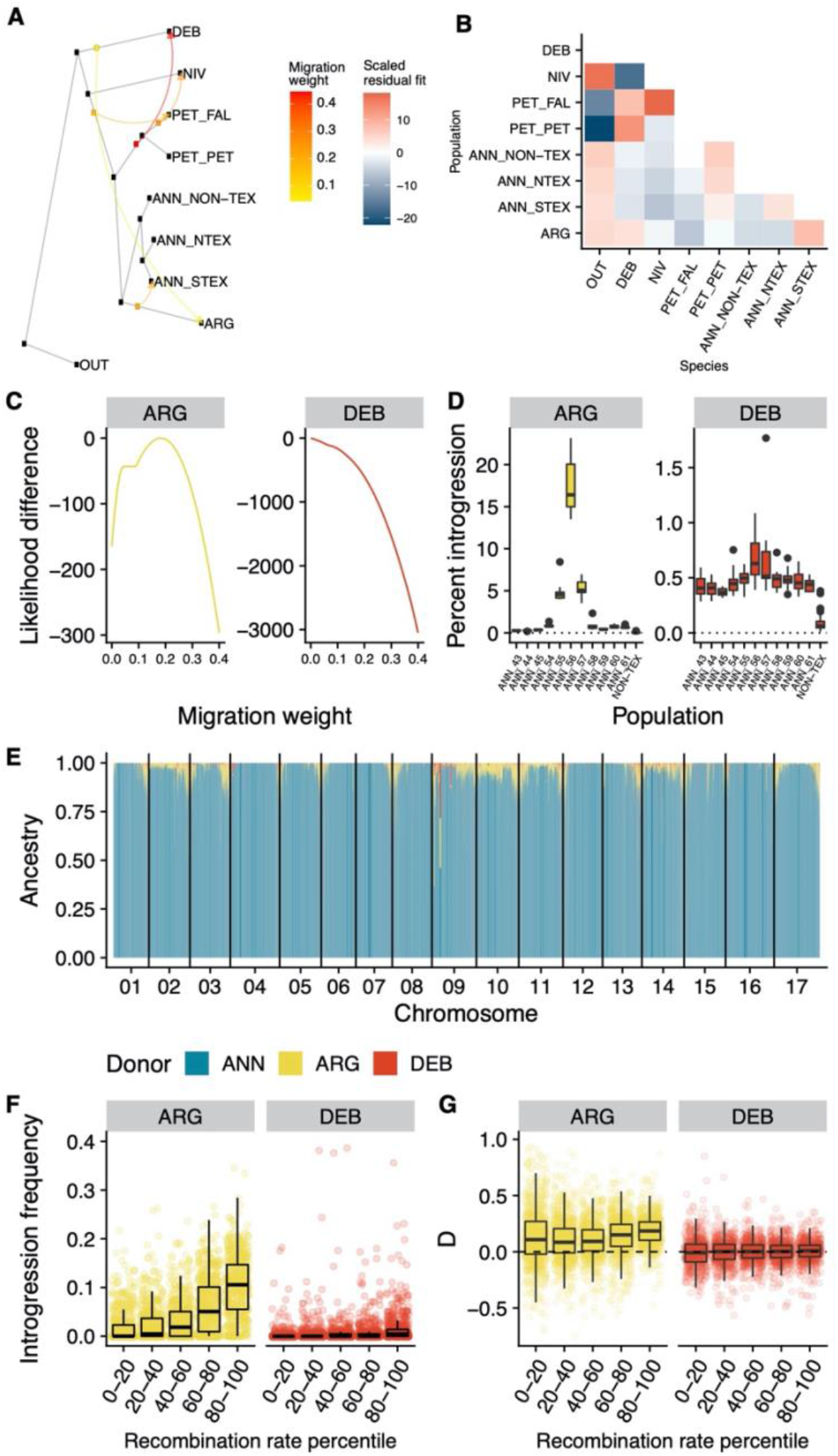
TreeMix and PCAdmix output testing for introgression. A. TreeMix output with 5 migration edges. B. Scaled residual fit after tree but without migration edges from TreeMix. C. Relative likelihood values for different weights of migration into H. a. texanus. D. Total percent introgression (>0.9 confidence) for H. annuus populations from PCAdmix. Non-Texas are H. a. annuus samples tested using a ‘leave one out’ strategy. E. Total percent introgressed ancestry for all H. a. texanus across the genome, scaled by basepair. F. Proportion introgressed ancestry for H. argophyllus (ARG) and H. debilis (DEB) in windows separated by recombination rate percentile. G. D in 100 SNP windows separated by recombination rate percentile for the donor species H. argophyllus (ARG) and H. debilis (DEB).

### Localized Introgression detection via Patterson’s D and PCAdmix

Across the genome we see much more variation and more positive values of D when *H. argophyllus* is used as a donor instead of *H. debilis* (Fig. S2). For *H. debilis*, there are two large regions with negative D scores which correspond to haploblocks on chromosome 5 and 13 (ann05.01, and ann13.02; Fig. S2) (Todesco et al., 2020).

At a total genome level, PCAdmix was consistent with D and showed high levels of *H. argophyllus* admixture in three populations of *H. a. texanus* (Fig. 4D). The inferred *H. debilis* introgression was much lower (~0.5%), although these values are higher than most, but not all, of the “control” non-Texas *H. a. annuus* samples. Across the genome, admixture was more common in the higher recombination regions on the ends of chromosomes, strongly for *H. argophyllus* and very slightly for *H. debilis* (Fig. 4E). We also found that D scores were higher for high recombination regions when *H. argophyllus* was the potential donor, but not when *H. debilis* was the donor (Fig. 4F).

Based on our methods, we also attempted to identify candidate introgressed regions. From *f_d_* measures testing for *H. debilis* gene flow into *H. a. texanus*, we selected the top 1% of windows, representing 101 regions spanning 49.3 Mbp. For PCAdmix, we required >40% *H. debilis* ancestry across all H. a. texanus samples, resulting in only 13 regions spanning 1.0 Mbp being selected (Fig. S5, File S2, S3). Between these two measures for detecting *H. debilis* introgression, we find two regions of overlap on chromosome 13 and 15. The only gene found in these regions is Ha412HOChr13g0625071 on chromosome 13. This gene shows similarity to anthocyanidin 3-O-glucosyltransferases, enzymes which are known to play a major role in the accumulation of anthocyanin pigments (Saito et al., 2013).

## Discussion

Heiser (1951a) hypothesized that morphologically divergent southeastern Texas forms of *Helianthus annuus* were the result of adaptive introgression from the narrowly distributed Texas sunflower *H. debilis*, with which they are morphologically convergent (Heiser, 1951a, Whitney et al., 2006, 2010). QTL mapping of *H. annuus* x *H. debilis* hybrids identified numerous candidate loci for which *H. debilis* alleles shift the phenotype towards *H. a. texanus*, as well as three candidate regions where *H. debilis* alleles are associated with increased fitness (Whitney et al., 2015). Further, field experimental evolution trials have shown that hybridization with *H. debilis* can allow populations to more quickly adapt over generations (Mitchell et al., 2019). This evidence has made *H. a. texanus* one of the few clear examples of adaptive introgression in plants (Suarez-Gonzalez et al., 2018).

We confirmed that *H. a. texanus* populations are indeed morphologically (and genetically) divergent from *H. a. annuus*, but four different approaches to detecting introgression found evidence for either very little (Admixture, Patterson’s D, PCAdmix) or no (TreeMix) introgression from *H. debilis* into *H. a. texanus*. Although *H. a. texanus* is genetically different from *H. a. annuus*, this difference does not seem to be due to introgression from *H. debilis*, but is concentrated in large segregating haploblocks (likely due to inversions), and is consistent with general isolation with distance patterns seen across the range (Todesco et al., 2020).

Despite this relatively unambiguous result, we have reasons to be cautious in our interpretation. Besides the morphological convergence that prompted the initial hypothesis, there are strong reasons to expect that *H. debilis* ancestry is present in *H. a. texanus*. Both species hybridize in the wild and *H. debilis* is the source of agronomically important traits in cultivated *H. annuus*, brought in through breeding (Jan & Chandler, 1985). Artificial introduction of *H. debilis* genetic material to *H. a. annuus*, followed by natural selection in the wild, can result in rapid fitness increases relative to control (non-introgressed) stock (Mitchell et al., 2019), suggesting that at least some introgressed *H. debilis* regions would be favored and retained in wild *H. a. texanus*. Thus, it is somewhat surprising that we detect so little admixture; below, we consider the evidence provided by each of the analyses we performed, and explore possible confounding factors masking the presence of *H. debilis* introgressions.

Patterson’s D is the most direct and model-free method for detecting introgression that we used, but can be confounded by introgression in other branches of the tree. Previous work has shown that *H. a. annuus* has gene flow with *H. petiolaris* (Kane et al., 2009), and our D scores using *H. petiolaris fallax* as the donor confirms that there is greater *H. petiolaris* introgression into the sympatric *H. a. annuus* than the allopatric *H. a. texanus*. Both *H. petiolaris* and *H. debilis* are within the same clade, so introgression from one species could be interpreted as introgression from the other (Fig. 3B). Thus a test of Patterson’s D using *H. a. annuus*, *H. a. texanus* and *H. debilis* is, to some extent, actually asking if there is greater gene flow between *H. a. texanus* and *H. debilis* versus between *H. a. annuus* and *H. petiolaris*. The degree that this would affect the test is hard to predict and depends on the divergence between *H. petiolaris* and *H. debilis*. To properly control for this, we would need a population of *H. a. annuus* without introgression; but given the broad range of *H. petiolaris* and likely continuous gene flow, we do not know of such a population.

This potential confounding factor is less relevant for TreeMix, which considers all species together, so if both gene flow events are occurring, TreeMix should detect them. TreeMix suggested several introgression edges that agreed with what is known in the system. For example, *H. petiolaris fallax* and *H. niveus canescens* overlap in range, intergrade together morphologically, and have high levels of introgression according to genotyping by sequencing analyses (Heiser et al., 1969; Zhang et al., 2019). Introgression from *H. argophyllus* into *H. a. texanus* and from *H. debilis* into *H. argophyllus* are both also supported by our ADMIXTURE results. Despite specifically testing for *H. debilis* into *H. a. texanus* introgression, TreeMix does not support any migration edge.

ADMIXTURE is similarly clear that *H. debilis* is not a part of *H. a. texanus* ancestry, with the exception of ANN1363. At K=3, ADMIXTURE does place both *H. debilis* and the *H. argophyllus* introgressed population ANN_56 in the same group, but this grouping is not consistent at other K values, or supported by other tests (Fig. S2). One of two equally-probable results from STRUCTURE is consistent with ADMIXTURE, while the other supports some *H. debilis* introgression in the same populations that have *H. argophyllus* introgression. In contrast with previous programs but partially consistent with one result from STRUCTURE, PCAdmix found that *H. a. texanus* samples had low but higher than baseline (allopatric *H. a. annuus*) levels of *H. debilis* and *H. argophyllus* ancestry (Fig. 4D). There are a few possible reasons why PCAdmix detected introgression not seen in other methods. PCAdmix is designed for recently admixed populations, so if *H. debilis* introgression is old, expectations of recombination in the PCAdmix model will likely be violated. Furthermore, PCAdmix is not robust to incomplete lineage sorting, which occurs at the high effective population sizes seen in sunflowers (Yuan et al., 2017). We attempted to use the allopatric *H. a. annuus* as a control but this is not perfect. *H. a. texanus* is genetically different from *H. a. annuus* (Fig. 2A), and this difference may drive *H. a. texanus* samples into intermediate positions in the two dimensional PCA, more so than a single non-Texas *H. a. annuus* sample, even in the absence of introgression.

Taken together, it’s unlikely that *H. debilis* contributed significantly to *H. a. texanus*’s ancestry, but that does not necessarily mean it did not contribute at all. In both D and PCAdmix, we found small regions that are consistent with *H. debilis* introgression into *H. a. texanus*, although we don’t know if they are due to incomplete lineage sorting (ILS) or introgression. Between PCAdmix and D we find two regions of overlap, which is perhaps not surprising considering both methods are looking for similar patterns. The single gene in those regions is involved in anthocyanidin production, and it is tempting to link this to the characteristic purple stems of *H. a. texanus,* but much further experimental work would be required to test this idea. Future work to QTL map herbivore resistance in *H. a. texanus*, a proposed adaptively introgressed trait, could cross reference with this list of regions to identify candidates, or rule out introgression as the source of the trait. It’s also possible that more complicated structural variation, e.g. copy number variation, is poorly captured in our SNP data and is introgressed but not detected in our scans.

How do we reconcile our paucity of introgression signals with previous genetic analyses finding the opposite? We suspect that new analyses methods not available to previous studies, like Patterson’s D, are better able to disentangle incomplete lineage sorting, population structure and introgression. For an in-depth overview of previous results, see the supplementary discussion.

In contrast with the minimal signal of introgression from *H. debilis*, there is unambiguous introgression from *H. argophyllus* into some *H. a. texanus* populations. We can see this in all except the most northern and western populations, which do not overlap with *H. argophyllus*’ distribution (Fig. 3D). The highest signals of introgression — up to 20% according to PCAdmix — are found in coastal populations of *H. a. texanus*. Interestingly, *H. argophyllus* presents two very distinct ecotypes (Moyers & Rieseberg, 2016). A late-flowering ecotype, which grows primarily inland, flowers later than most *H. annuus* populations, while an early-flowering ecotype’s flowering time overlaps with *H. annuus*. The early-flowering ecotype occurs on the coast, where signals of introgression are strongest. This suggests that flowering time is an important reproductive barrier between these species.

The introgression from *H. argophyllus* likely contains deleterious incompatibility loci that are being selected against. We see this in the positive relationship between introgression and recombination rate (Fig. 4F-G). This pattern can arise because higher recombination allows for deleterious incompatibility loci to be decoupled from the introgressed haplotype more rapidly (Nachman & Payseur, 2012; Schumer et al., 2018). That being said, we also see a negative relationship between F_ST_ and recombination rate even when populations are allopatric (Fig. 2E-F) which supports a role for linked selection (Burri, 2017). Given that sunflowers have large regions of extremely low recombination, polymorphic inversions and rampant structural changes between species, it will be important but challenging to account the effects of recombination when comparing genomes between species (Todesco et al., 2020; Ostevik et al., 2020).

## Conclusion

The lack of a clear signal of introgression from *H. debilis* into *H. a. texanus* suggests that Heiser’s (1951a) original hypothesis needs to be reassessed. There are three possible interpretations for the results we present. First, if we make the assumption that the trait convergence must come from introgression, then our results suggest that the minimal *H. debilis* introgression contains important alleles. Some of our results suggest that *H. argophyllus* could have acted as a bridge species between *H. debilis* and *H. annuus*; in our Patterson’s D test, the highest signals of *H. debilis* introgression in Texas *H. annuus* (excluding ANN1363) occur in samples with high *H. argophyllus* introgression and there is evidence of introgression from *H. debilis* into *H. argophyllus*. However, since *H. argophyllus* does not show phenotypic convergence with *H. a. texanus* and *H. debilis*, it seems unlikely that it would have facilitated sharing of alleles responsible for the convergence between these two species.

Alternatively, we could reassess the entire premise of historic adaptive introgression in this system. Although *H. a. texanus* is statistically different from *H. a. annuus* at several traits, its values almost never exceed the range of values seen in *H. a. annuus*. It seems therefore possible that the morphology of *H. a. texanus* could have evolved from standing variation and does not require introgression.

Finally, it is also possible that introgression has played a role in some, but not all, the traits that define *H. a. texanus*. To truly know whether adaptive introgression has occurred in a hybrid lineage, the alleles underlying the adaptive trait need to be identified and phylogenetically compared with possible donor species. Here we have identified a potential introgressed candidate gene for pigmentation differences between Texas and non-Texas *H. annuus*. Follow-up work, looking at genetic and functional diversity at this locus in *H. annuus* and *H. debilis*, will be required to confirm its introgressed origin and its role in the convergence between these two species.

## Acknowledgements

Funding was provided by Genome Canada and Genome BC (LSARP2014-223SUN), the National Science Foundation (NSF DEB 1257965) and a Banting postdoctoral fellowship to G.L.O.

## Author Contributions

G.L.O., N.M., K.D.W. and L.H.R. conceived of the project. M.T. and N.B. collected genomic and phenotypic data. G.L.O. and J.S.M. processed sequence data. G.L.O. conducted all analyses.

G.L.O. wrote the manuscript with contributions from all authors.

## Data Accessibility

All raw sequence data is available on the Sequence Read Archive (SRA). Identification numbers are listed in supplementary file 1.

## References

Abbott R, Albach D, Ansell S, Arntzen JW, Baird SJ, Bierne N, Boughman J, Brelsford A, Buerkle CA, Buggs R, Butlin RK. Hybridization and speciation. Journal of evolutionary biology. 2013 Feb;26(2):229–46.

Alexander DH, Lange K. Enhancements to the ADMIXTURE algorithm for individual ancestry estimation. BMC bioinformatics. 2011 Dec 1;12(1):246.

Arnold ML, Kunte K. Adaptive genetic exchange: a tangled history of admixture and evolutionary innovation. Trends in ecology & evolution. 2017 Aug 1;32(8):601–11.

Badouin H, Gouzy J, Grassa CJ, Murat F, Staton SE, Cottret L, Lelandais-Brière C, Owens GL, Carrère S, Mayjonade B, Legrand L. The sunflower genome provides insights into oil metabolism, flowering and Asterid evolution. Nature. 2017 Jun;546(7656):148–52.

Bolger AM, Lohse M, Usadel B. Trimmomatic: a flexible trimmer for Illumina sequence data. Bioinformatics. 2014 Aug 1;30(15):2114–20.

Brisbin A, Bryc K, Byrnes J, Zakharia F, Omberg L, Degenhardt J, Reynolds A, Ostrer H, Mezey JG, Bustamante CD. PCAdmix: principal components-based assignment of ancestry along each chromosome in individuals with admixed ancestry from two or more populations. Human biology. 2012 Aug;84(4):343.

Browning SR, Browning BL. Rapid and accurate haplotype phasing and missing-data inference for whole-genome association studies by use of localized haplotype clustering. The American Journal of Human Genetics. 2007 Nov 1;81(5):1084–97.

Burri R. Interpreting differentiation landscapes in the light of long‐term linked selection. Evolution Letters. 2017 Aug;1(3):118–31.

Chandler JM, Jan CC, Beard BH. Chromosomal differentiation among the annual *Helianthus species*. Systematic Botany. 1986 Apr 1:354–71

Giska I, Farelo L, Pimenta J, Seixas FA, Ferreira MS, Marques JP, Miranda I, Letty J, Jenny H, Hackländer K, Magnussen E. Introgression drives repeated evolution of winter coat color polymorphism in hares. Proceedings of the National Academy of Sciences. 2019 Nov 26;116(48):24150–6.

Harrison RG, Larson EL. Hybridization, introgression, and the nature of species boundaries. Journal of Heredity. 2014 Dec 1;105(S1):795–809.

Heiser Jr CB. Hybridization between the sunflower species *Helianthus annuus* and *H. petiolaris*. Evolution. 1947 Dec;1(4):249–62.

Heiser Jr CB. Study in the evolution of the sunflower species *Helianthus annuus* and *H. bolanderi*. Botany. 1949;23:157–208.

Heiser Jr CB. Hybridization in the annual sunflowers: *Helianthus annuus* × *H. debilis var. cucumerifolius*. Evolution. 1951a Mar 1:42–51.

Heiser Jr CB. Hybridization in the annual sunflowers: *Helianthus annuus* x H. argophyllus. The American Naturalist. 1951b Jan 1;85(820):65–72.

Heiser Jr CB. Variation and subspeciation in the common sunflower, *Helianthus annuus*. American Midland Naturalist. 1954 Jan 1:287–305.

Heiser Jr CB, Smith DM, Clevenger SB, Martin WC. The north american sunflowers (*Helianthus*). Memoirs of the Torrey Botanical Club. 1969 Jan 31;22(3):1–218.

Hovick, S. M. & Whitney, K. D. Hybridisation is associated with increased fecundity and size in invasive taxa: meta-analytic support for the hybridisation-invasion hypothesis. Ecology Letters 17, 1464–1477 (2014).

Jan CC, Chandler JM. Transfer of powdery mildew resistance from Helianthus debilis Nutt. To cultivated sunflower 1. Crop Science. 1985;25(4):664–6.

Jay P, Whibley A, Frézal L, de Cara MÁ, Nowell RW, Mallet J, Dasmahapatra KK, Joron M. Supergene evolution triggered by the introgression of a chromosomal inversion. Current Biology. 2018 Jun 4;28(11):1839–45.

Jones MR, Mills LS, Alves PC, Callahan CM, Alves JM, Lafferty DJ, Jiggins FM, Jensen JD, Melo-Ferreira J, Good JM. Adaptive introgression underlies polymorphic seasonal camouflage in snowshoe hares. Science. 2018 Jun 22;360(6395):1355–8.

Josse J, Husson F. missMDA: a package for handling missing values in multivariate data analysis. Journal of Statistical Software. 2016 Apr 4;70(1):1–31.

Kane NC, King MG, Barker MS, Raduski A, Karrenberg S, Yatabe Y, Knapp SJ, Rieseberg LH. Comparative genomic and population genetic analyses indicate highly porous genomes and high levels of gene flow between divergent *Helianthus species*. Evolution: International Journal of Organic Evolution. 2009 Aug;63(8):2061–75.

Kopelman NM, Mayzel J, Jakobsson M, Rosenberg NA, Mayrose I. Clumpak: a program for identifying clustering modes and packaging population structure inferences across K. Molecular ecology resources. 2015 Sep;15(5):1179–91.

Lawlor J. jakelawlor/PNWColors: Initial Release. Zenodo; 2020.

Lê S, Josse J, Husson F. FactoMineR: an R package for multivariate analysis. Journal of statistical software. 2008 Mar 18;25(1):1–8.

Li H. A statistical framework for SNP calling, mutation discovery, association mapping and population genetical parameter estimation from sequencing data. Bioinformatics. 2011 Nov 1;27(21):2987–93.

Martin SH, Davey JW, Jiggins CD. Evaluating the use of ABBA-BABA statistics to locate introgressed loci. Molecular biology and evolution. 2015 Jan 1;32(1):244–57.

Mitchell N, Owens GL, Hovick SM, Rieseberg LH, Whitney KD. Hybridization speeds adaptive evolution in an eight-year field experiment. Scientific reports. 2019 May 1;9(1):1–2.

Moyers BT, Rieseberg LH. Remarkable life history polymorphism may be evolving under divergent selection in the silverleaf sunflower. Molecular Ecology. 2016 Aug;25(16):3817–30.

Nachman MW, Payseur BA. Recombination rate variation and speciation: theoretical predictions and empirical results from rabbits and mice. Philosophical Transactions of the Royal Society B: Biological Sciences. 2012 Feb 5;367(1587):409–21.

Ostevik KL, Samuk K, Rieseberg LH. Ancestral Reconstruction of Karyotypes Reveals an Exceptional Rate of Nonrandom Chromosomal Evolution in Sunflower. Genetics. 2020 Apr 1;214(4):1031–45.

Owens GL, Rieseberg LH. Hybrid incompatibility is acquired faster in annual than in perennial species of sunflower and tarweed. Evolution. 2014 Mar;68(3):893–900.

Owens GL, Baute GJ, Rieseberg LH. Revisiting a classic case of introgression: hybridization and gene flow in Californian sunflowers. Molecular ecology. 2016 Jun;25(11):2630–43.

Patterson N, Moorjani P, Luo Y, Mallick S, Rohland N, Zhan Y, Genschoreck T, Webster T, Reich D. Ancient admixture in human history. Genetics. 2012 Nov 1;192(3):1065–93.

Petr M, Vernot B, Kelso J. admixr—R package for reproducible analyses using ADMIXTOOLS. Bioinformatics. 2019 Sep 1;35(17):3194–5.

Pickrell J, Pritchard J. Inference of population splits and mixtures from genome-wide allele frequency data. Nature Precedings. 2012

Pritchard JK, Stephens M, Donnelly P. Inference of population structure using multilocus genotype data. Genetics. 2000 Jun 1;155(2):945–59.

Purcell S, Neale B, Todd-Brown K, Thomas L, Ferreira MA, Bender D, Maller J, Sklar P, De Bakker PI, Daly MJ, Sham PC. PLINK: a tool set for whole-genome association and population-based linkage analyses. The American journal of human genetics. 2007 Sep 1;81(3):559–75.

R Core Team. R: A language and environment for statistical computing.

Rieseberg, L. H. 1991. Homoploid reticulate evolution in *Helianthus* (Asteraceae): evidence from ribosomal genes. American Journal of Botany 78:1218–1237.

Rieseberg LH, Beckstrom-Sternberg S, Doan K. *Helianthus annuus ssp. texanus* has chloroplast DNA and nuclear ribosomal RNA genes of *Helianthus debilis ssp*. cucumerifolius. Proceedings of the National Academy of Sciences. 1990a Jan 1;87(2):593–7.

Rieseberg LH, Carter R, Zona S. Molecular tests of the hypothesized hybrid origin of two diploid *Helianthus species* (Asteraceae). Evolution. 1990b Sep;44(6):1498–511.

Rieseberg LH, Kim SC, Randell RA, Whitney KD, Gross BL, Lexer C, Clay K. Hybridization and the colonization of novel habitats by annual sunflowers. Genetica. 2007 Feb 1;129(2):149–65.

Saito K, Yonekura-Sakakibara K, Nakabayashi R, Higashi Y, Yamazaki M, Tohge T, Fernie AR. The flavonoid biosynthetic pathway in Arabidopsis: structural and genetic diversity. Plant Physiol Biochem. 2013 Nov;72:21–34

Scascitelli M, Whitney KD, Randell RA, King M, Buerkle CA, Rieseberg LH. Genome scan of hybridizing sunflowers from Texas (*Helianthus annuus* and *H. debilis*) reveals asymmetric patterns of introgression and small islands of genomic differentiation. Molecular Ecology. 2010 Feb;19(3):521–41.

Schumer M, Xu C, Powell DL, Durvasula A, Skov L, Holland C, Blazier JC, Sankararaman S, Andolfatto P, Rosenthal GG, Przeworski M. Natural selection interacts with recombination to shape the evolution of hybrid genomes. Science. 2018 May 11;360(6389):656–60.

Sedlazeck FJ, Rescheneder P, Von Haeseler A. NextGenMap: fast and accurate read mapping in highly polymorphic genomes. Bioinformatics. 2013 Nov 1;29(21):2790–1.

Seehausen O. Hybridization and adaptive radiation. Trends in ecology & evolution. 2004 Apr 1;19(4):198–207.

Stephens JD, Rogers WL, Mason CM, Donovan LA, Malmberg RL. Species tree estimation of diploid Helianthus (Asteraceae) using target enrichment. American Journal of Botany. 2015 Jun;102(6):910–20.

Suarez-Gonzalez A, Lexer C, Cronk QC. Adaptive introgression: a plant perspective. Biology letters. 2018 Mar 31;14(3):20170688.

[dataset] Todesco M, Owens GL, Bercovich N; 2020; Wild Helianthus GWAS and GEA; SRA: PRJNA532579

Todesco M, Owens GL, Bercovich N, Légaré JS, Soudi S, Burge DO, Huang K, Ostevik KL, Drummond EB, Imerovski I, Lande K, Pascual-Robles MA, Nanavati M, Jahani M, Cheung W, Staton SE, Muños S, Nielsen R, Donovan LA, Burke JM, Yeaman S, Rieseberg LH. Massive haplotypes underlie ecotypic differentiation in sunflowers. Nature. 2020 Aug;584(7822):602–7.

Van der Auwera GA, Carneiro MO, Hartl C, Poplin R, Del Angel G, Levy‐Moonshine A, Jordan T, Shakir K, Roazen D, Thibault J, Banks E. From FastQ data to high‐confidence variant calls: the genome analysis toolkit best practices pipeline. Current protocols in bioinformatics. 2013 Oct;43(1):11–0.

Weir BS, Cockerham CC. Estimating F‐statistics for the analysis of population structure. evolution. 1984 Nov;38(6):1358–70

Whitney KD, Randell RA, Rieseberg LH. Adaptive introgression of herbivore resistance traits in the weedy sunflower *Helianthus annuus*. The American Naturalist. 2006 Jun;167(6):794–807.

Whitney KD, Randell RA, Rieseberg LH. Adaptive introgression of abiotic tolerance traits in the sunflower *Helianthus annuus*. New Phytologist. 2010 Jul;187(1):230–9.

Whitney, K. D., K. W. Broman, N. C. Kane, S. M. Hovick, R. A. Randell, and L. H. Rieseberg. 2015. QTL mapping identifies candidate alleles involved in adaptive introgression and range expansion in a wild sunflower. Molecular Ecology 24: 2194–2211.

Wickham H, Averick M, Bryan J, Chang W, McGowan LD, François R, Grolemund G, Hayes A, Henry L, Hester J, Kuhn M. Welcome to the Tidyverse. Journal of Open Source Software. 2019 Nov 21;4(43):1686.

Yatabe Y, Kane NC, Scotti-Saintagne C, Rieseberg LH. Rampant gene exchange across a strong reproductive barrier between the annual sunflowers, *Helianthus annuus* and H. petiolaris. Genetics. 2007 Apr 1;175(4):1883–93.

Yuan K, Zhou Y, Ni X, Wang Y, Liu C, Xu S. Models, methods and tools for ancestry inference and admixture analysis. Quantitative Biology. 2017 Sep 1;5(3):236–50.

Zhang JQ, Imerovski I, Borkowski K, Huang K, Burge D, Rieseberg LH. Intraspecific genetic divergence within *Helianthus niveus* and the status of two new morphotypes from Mexico. American journal of botany. 2019 Sep;106(9):1229–39.

Zheng X, Levine D, Shen J, Gogarten SM, Laurie C, Weir BS. A high-performance computing toolset for relatedness and principal component analysis of SNP data. Bioinformatics. 2012 Dec 1;28(24):3326–8.

