## Supplementary Information for "An unexpected donor in the adaptive introgression candidate *Helianthus annuus* subsp. *texanus*"

#### Supplementary Discussion

How do we explain previous studies that suggested introgression from *H. debilis* into *H. a. texanus*? The first work to show this used cpDNA and rDNA restriction digests (Rieseberg et al. 1990a). They found a single cpDNA pattern in non-Texas *H. annuus*, while *H. debilis* had roughly a split between a unique cpDNA pattern and the non-Texas *H. annuus* pattern. In *H. a. texanus*, 6% of the samples had the *H. debilis* pattern, while the rest had matched the *H. a. annuus* pattern. Recent work using next-generation sequencing (NGS) found two main cpDNA lineages across the annual sunflowers (Lee-Yaw et al. 2019). Both *H. annuus* and *H. debilis* primarily contain cpDNA from Clade II, although they each have samples containing Clade I cpDNA. It's unclear whether the cpDNA restriction patterns represent the two clades seen in NGS data, or variation within the clades.

The examination of nuclear markers (ribosomal DNA, rDNA) found some *H. debilis* markers in 10% of *texanus* samples (Riesberg et al. 1990a). While this may represent introgression, it could also be a product of ILS. Although this study used markers that were fixed for alternate alleles in reference panels, those panels were of limited size (30 and 16) so it is possible that the markers were not actually fixed differences, but polymorphic within *H. annuus*. This chance increases in the presence of population structure, which we now know exists between non-Texas and Texas *H. annuus* (Todesco et al. 2020). To test this, we subsampled 30 random non-Texas *H. a. annuus* samples and selected all sites with fixed differences between this subset and all eight sequenced *H. debilis*. We then measured the allele frequency in *H. a. texanus*. We found that for these markers that are ostensibly fixed differences, 36% of loci were at >1% minor allele frequency in *H. a. texanus*, while 4% were at >10% minor allele frequency. Thus, isolated introgression-like signals can be found in a dataset with no overall pattern of introgression.

A follow up study found higher levels of cpDNA discordance, but slightly lower levels of rDNA discordance (7%) (Rieseberg et al. 2007). At the time, ILS was dismissed because *H. annuus* and *H. debilis* are in separate, divergent clades. In our current filtered dataset, non-Texas *H. annuus* and *H. debilis* share polymorphism at 18% of loci, suggesting much higher amounts of shared variation than previously appreciated.

A more recent study (Scascitelli et al. 2010) used 88 microsatellite loci and found, again, very low levels of possible introgression. Using STRUCTURE, the authors found 3/150 *H. a. texanus* samples were early generation admixed with *H. debilis*, while 1/90 *H. a. annuus* samples were similarly admixed. No other *H. annuus* samples had admixture levels significantly above zero. Although this study estimated a non-zero migration rate from *H. debilis* into *H. a.*

*texanus*, it estimated a similar rate into *H. a. annuus*, which is geographically implausible, suggesting it did not accurately capture recent migration rates.

All together, previous work found low and inconsistent *H. debilis* ancestry in *H. a. texanus*. Although this has consistently been interpreted that *H. a. texanus* is a hybrid lineage, with more complete genomic data it seems like this is more likely due to ILS or possibly introgression through *H. argophyllus*.

### Supplementary Figures:

**Figure S1**: Trait comparisons for all traits thought to differentiate *H. a. annuus* and *H. a. texanus*.

P-values are Wilcox rank sum tests against Non-Texas *H. annuus*. For axis label “pos” means *H. a. texanus* expected to have a higher value and “neg” means it is expected to have a lower value.

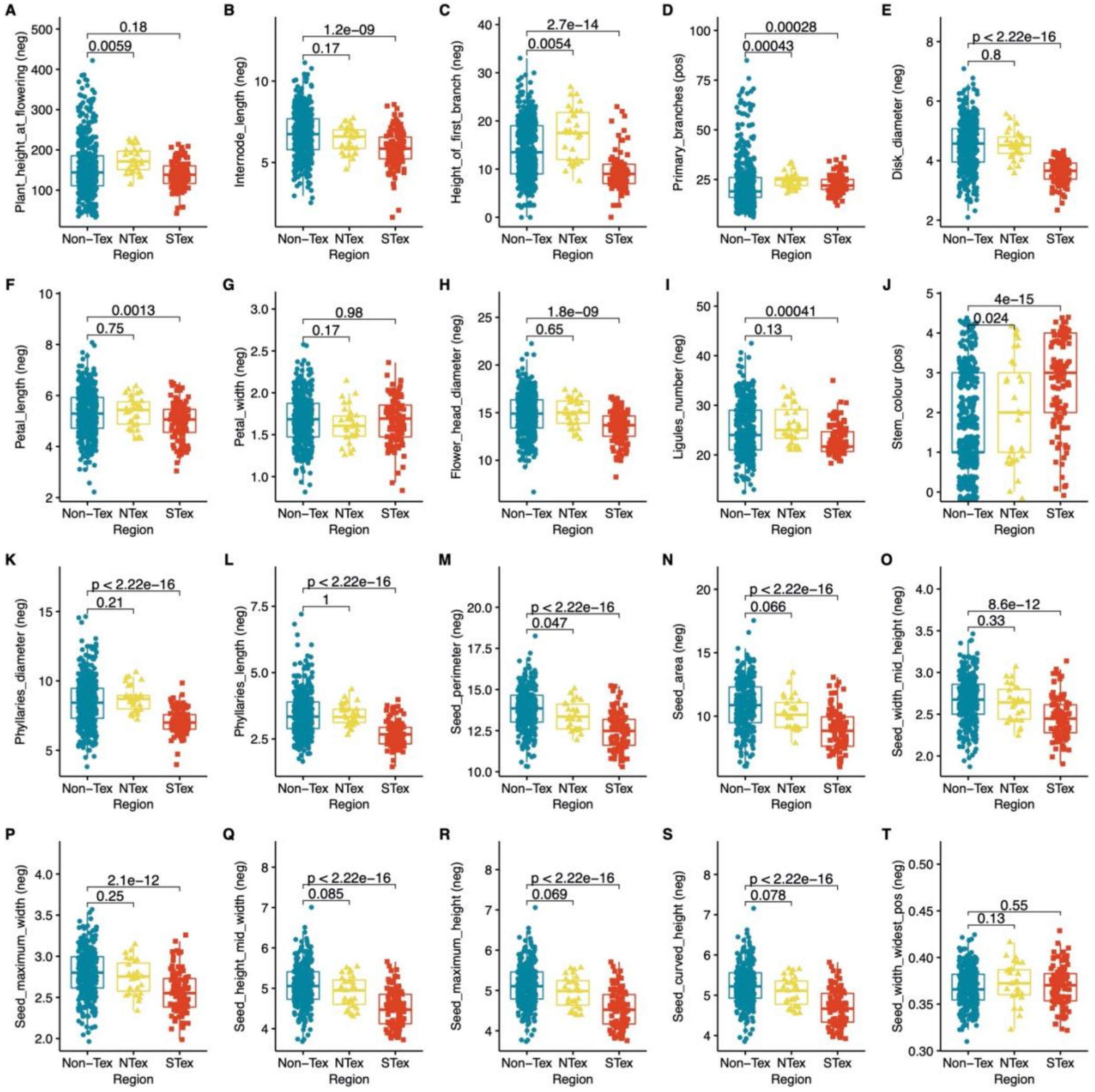

**Figure S2**: Genome wide D statistic.

A. D in 100 SNP windows using the arrangement *H. a. annuus*, *H. a. texanus*, *H. debilis*, outgroup. Segregating haploblocks in *H. annuus* showing signals different from background are highlighted. B. D in 100 SNP windows using the arrangement *H. a. annuus*, *H. a. texanus*, *H. argophyllus*, outgroup. C. D in 100 SNP windows using the arrangement *H. a. annuus*, ANN1363, *H. debilis*, outgroup. This sample has a clear *H. debilis* introgression signal on chromosome 3.

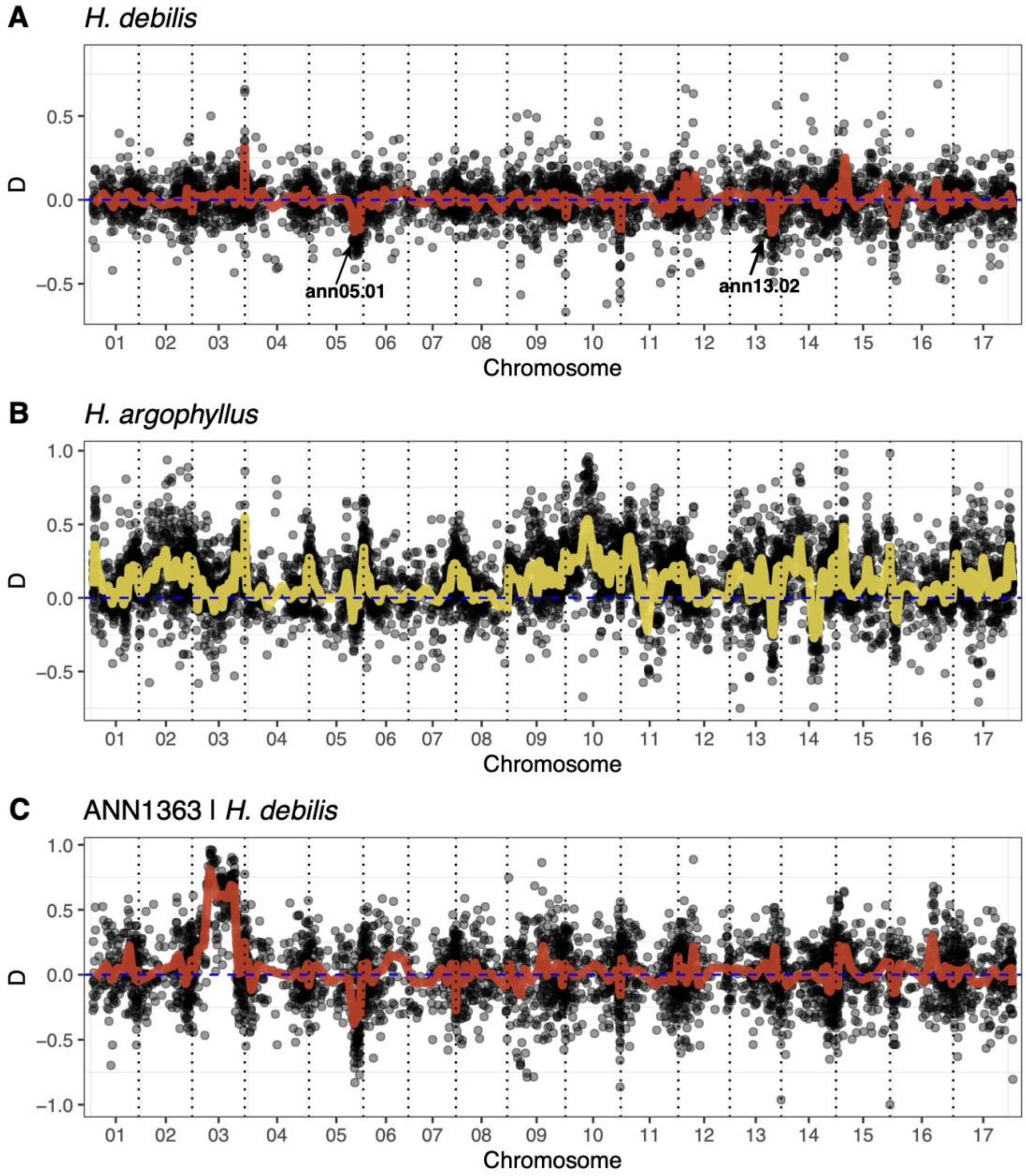

**Figure S3**: ADMIXTURE results for K=2 to K=10.

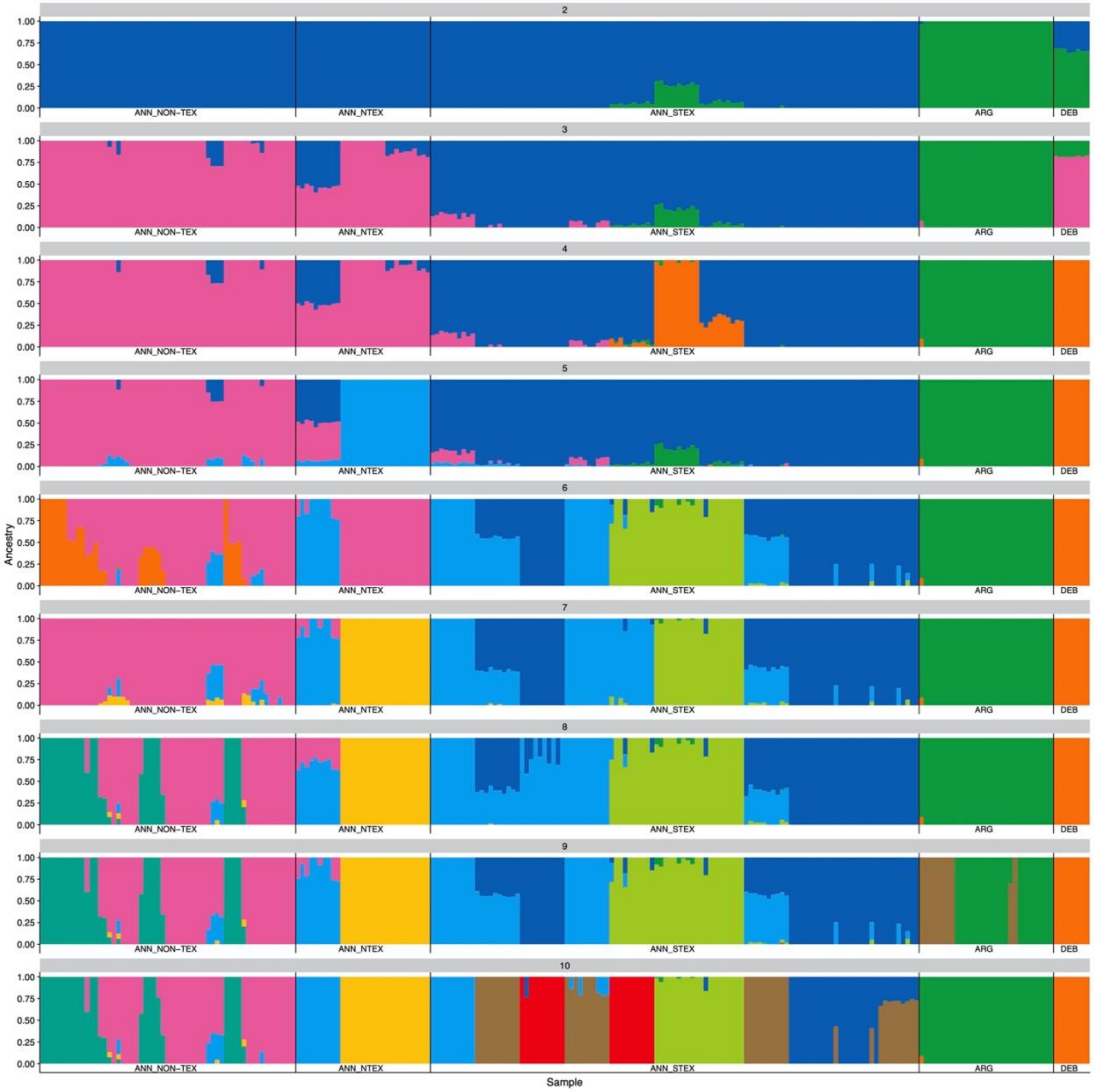

**Figure S4**: All treemix results. A-K. Tree and migration edges for 0 to 10 migration edges. L. Likelihood change for adding an additional migration edge from 1 to 10.

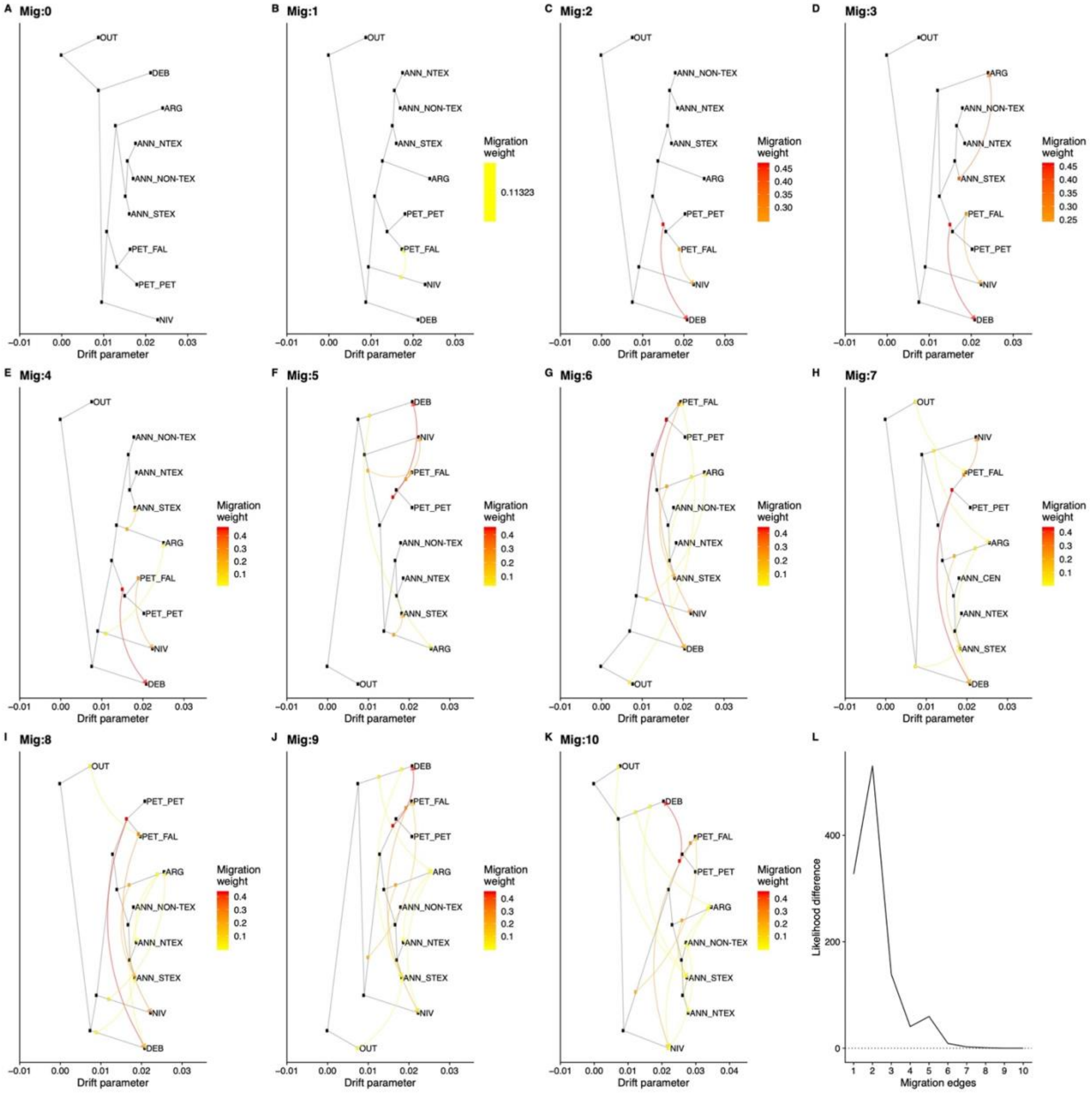

**Figure S5**: D and PCAdmix introgression regions, as well as overlapping regions. Overlapping regions are highlighted in yellow. Markers are not sized to scale.

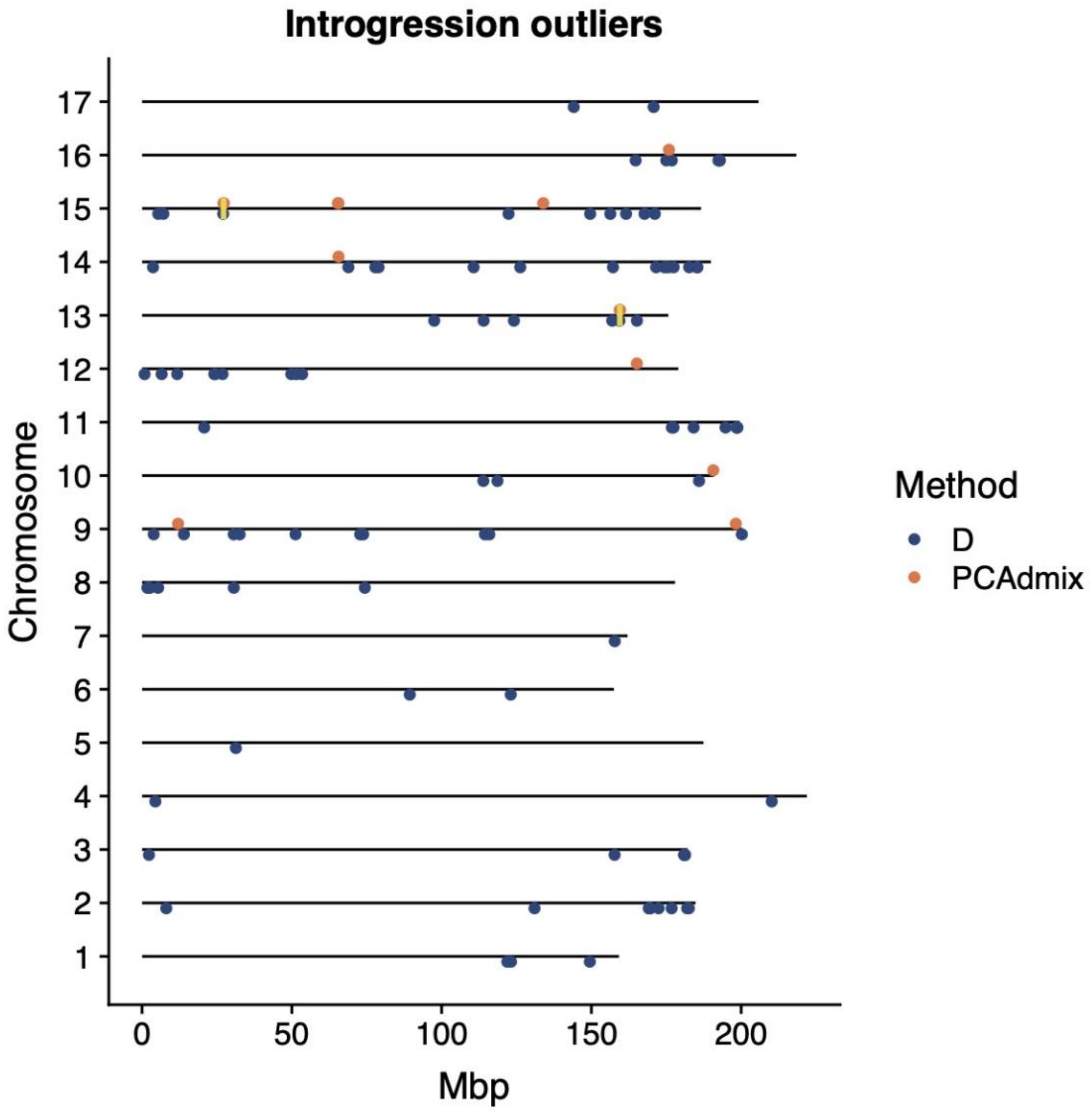

#### Supplementary Files:

**File S1:** Sample information.
Species, population, location and Sequence Read Archive number for all samples used.
**File S2**: *H. debilis* introgression candidates from D.

The top 2% of 100 window wide windows for $f_d$ using the set [*H. a. annuus*, *H. a. texanus*, *H. debilis*, perennials]

**File S3**: *H. debilis* introgression candidates from PCAdmix.

All windows with >40% *H. debilis* ancestry across all *H. a. texanus* samples.

#### References:


Lee-Yaw JA, Grassa CJ, Joly S, Andrew RL, Rieseberg LH. An evaluation of alternative explanations for widespread cytonuclear discordance in annual sunflowers (*Helianthus*). New Phytologist. 2019 Jan;221(1):515-26.

Rieseberg LH, Beckstrom-Sternberg S, Doan K. *Helianthus annuus ssp. texanus* has chloroplast DNA and nuclear ribosomal RNA genes of *Helianthus debilis ssp. cucumerifolius*. Proceedings of the National Academy of Sciences. 1990a Jan 1;87(2):593-7.

Rieseberg LH, Kim SC, Randell RA, Whitney KD, Gross BL, Lexer C, Clay K. Hybridization and the colonization of novel habitats by annual sunflowers. Genetica. 2007 Feb 1;129(2):149-65.

Scascitelli M, Whitney KD, Randell RA, King M, Buerkle CA, Rieseberg LH. Genome scan of hybridizing sunflowers from Texas (*Helianthus annuus* and *H. debilis*) reveals asymmetric patterns of introgression and small islands of genomic differentiation. Molecular Ecology. 2010 Feb;19(3):521-41.

Todesco M, Owens GL, Bercovich N, Légaré JS, Soudi S, Burge DO, Huang K, Ostevik KL, Drummond EB, Imerovski I, Lande K, Pascual-Robles MA, Nanavati M, Jahani M, Cheung W, Staton SE, Muños S, Nielsen R, Donovan LA, Burke JM, Yeaman S, Rieseberg LH. Massive haplotypes underlie ecotypic differentiation in sunflowers. Nature. 2020 Aug;584(7822):602-7.
